# Genetic validation of bipolar disorder identified by automated phenotyping using electronic health records

**DOI:** 10.1101/193011

**Authors:** Chia-Yen Chen, Phil H. Lee, Victor M. Castro, Jessica Minnier, Alexander W. Charney, Eli A. Stahl, Douglas M. Ruderfer, Shawn N. Murphy, Vivian Gainer, Tianxi Cai, Ian Jones, Carlos Pato, Michele Pato, Mikael Landén, Pamela Sklar, Roy H. Perlis, Jordan W. Smoller

**Affiliations:** Psychiatric and Neurodevelopmental Genetics Unit, Massachusetts General Hospital, 185 Cambridge St., Boston, MA 02114, USA; Analytic and Translational Genetics Unit, Center for Human Genetic Research, Massachusetts General Hospital, 185 Cambridge St., Boston, MA 02114, USA; Center for Genomic Medicine, Massachusetts General Hospital, 185 Cambridge St, Boston, MA 02114, USA.; Department of Psychiatry, Massachusetts General Hospital, 55 Fruit Street, Boston, MA 02114, USA; Broad Institute of MIT and Harvard, 75 Ames Street, Cambridge, MA 02142, USA.; Center for Experimental Drugs and Diagnostics, Massachusetts General Hospital, 55 Fruit Street, Boston, MA 02114, USA; Partners Research Information Systems and Computing, Partners HealthCare System, One Constitution Center, Charlestown, MA 02129, USA; Oregon Health & Sciences University, 3181 SW Sam Jackson Park Rd, Portland, OR 97239, USA; Department of Psychiatry, Icahn School of Medicine at Mount Sinai, One Gustave L. Levy Place, New York, NY 10029, USA; Institute for Genomics and Multiscale Biology, Department of Genetics and Genomic Sciences, Icahn School of Medicine at Mount Sinai, One Gustave L. Levy Place, New York, NY 10029, USA; Friedman Brain Institute, Department of Neuroscience, Icahn School of Medicine at Mount Sinai, One Gustave L. Levy Place, New York, NY 10029, USA; Division of Genetic Medicine, Vanderbilt University Medical Center, Nashville, TN 37212, USA; Department of Neurology, Massachusetts General Hospital, 55 Fruit Street, Boston, MA 02114, USA; Department of Biomedical Informatics, Harvard Medical School, 10 Shattuck Street, Boston, MA 02115, USA; Department of Biostatistics, Harvard T.H. Chan School of Public Health, 677 Huntington Ave, Boston, MA 02115, USA; National Centre for Mental Health, MRC Centre for Neuropsychiatric Genetics and Genomics, Cardiff University, Cardiff, CF24 4HQ, UK; SUNY Downstate Medical Center, Brooklyn, NY 11203, USA; Institute of Neuroscience and Physiology, Department of Psychiatry and Neurochemistry, The Sahlgrenska Academy, University of Gothenburg, Gothenburg, Sweden; Department of Medical Epidemiology and Biostatistics, Karolinska Institutet, Stockholm, Sweden

**Keywords:** Bipolar disorder, electronic health records, phenotyping, genetic, heritability

## Abstract

Bipolar disorder (BD) is a heritable mood disorder characterized by episodes of mania and depression. Although genomewide association studies (GWAS) have successfully identified genetic loci contributing to BD risk, sample size has become a rate-limiting obstacle to genetic discovery. Electronic health records (EHRs) represent a vast but relatively untapped resource for high-throughput phenotyping. As part of the International Cohort Collection for Bipolar Disorder (ICCBD), we previously validated automated EHR-based phenotyping algorithms for BD against in-person diagnostic interviews (Castro et al. 2015). Here, we establish the genetic validity of these phenotypes by determining their genetic correlation with traditionally-ascertained samples. Case and control algorithms were derived from structured and narrative text in the Partners Healthcare system comprising more than 4.6 million patients over 20 years. Genomewide genotype data for 3,330 BD cases and 3,952 controls of European ancestry were used to estimate SNP-based heritability (h^2^_g_) and genetic correlation(r_g_) between EHR-based phenotype definitions and traditionally-ascertained BD cases in GWAS by the ICCBD and Psychiatric Genomics Consortium (PGC) using LD score regression. We evaluated BD cases identified using 4 EHR-based algorithms: an NLP-based algorithm (95-NLP) and 3 rule-based algorithms using codified EHR with decreasing levels of stringency - “coded-strict”, “coded-broad”, and “coded-broad based on a single clinical encounter” (coded-broad-SV). The analytic sample comprised 862 95-NLP, 1,968 coded-strict, 2,581 coded-broad, 408 coded-broad-SV BD cases, and 3,952 controls. The estimated h^2^_g_ were 0.24 (p=0.015), 0.09 (p=0.064), 0.13 (p=0.003), 0.00 (p=0.591) for 95-NLP, coded-strict, coded-broad and coded-broad-SV BD, respectively. The h^2^_g_ for all EHR-based cases combined except coded-broad-SV (excluded due to 0 h^2^_g_) was 0.12 (p=0.004). These h^2^_g_ were lower or similar to the h^2^_g_ observed by the ICCBD+PGCBD (0.23, p=3.17E-80, total N=33,181). However, the r_g_ between ICCBD+PGCBD and the EHR-based cases were high for 95-NLP (0.66, p=3.69x10-5), coded-strict (1.00, p=2.40x10-4), and coded-broad (0.74, p=8.11x10-7). The r_g_ between EHR-based BDs ranged from 0.90 to 0.98. These results provide the first genetic validation of automated EHR-based phenotyping for BD and suggest that this approach identifies cases that are highly genetically correlated with those ascertained through conventional methods. High throughput phenotyping using the large data resources available in EHRs represents a viable method for accelerating psychiatric genetic research.

## Introduction

Although twin studies first documented the high heritability of bipolar disorder (BD) decades ago, only recently have robustly associated genetic risk loci been identified through genomewide association studies (GWAS).^1-8^ At present, the major rate-limiting step for GWAS of BD is the need for ever-larger sample sizes to detect both common modest-effect variants and rarer large effect variants. In recent years, the widespread adoption of longitudinal electronic health records (EHRs) has provided a vast and growing repository of phenotypic data that can be leveraged for psychiatric research.^9^ In particular, when linked to sample collections through biobanks and other efforts, EHR data provide a relatively untapped opportunity to enhance the power of genetic research. Nevertheless, establishing the validity of EHR-derived phenotypes remains an important pre-requisite for leveraging these resources.

In an effort to rapidly increase available samples for genomewide studies of BD, we established the International Cohort Collection for Bipolar Disorder (ICCBD) through which we applied high-throughput phenotyping methods at sites in the United States (US), United Kingdom (UK) and Sweden.^7^ At the US site (Partners Healthcare), we developed and applied EHR phenotyping algorithms to identify approximately 4,500 cases and 5,000 controls for whom DNA was obtained from discarded blood samples. The use of EHR data to define valid phenotypes is particularly challenging for psychiatric disorders. Because there are no pathognomonic laboratory or pathologic findings, psychiatric diagnosis has traditionally relied on self-reported symptoms, behavioral observations, and clinical judgment. Thus, genomic studies have typically utilized structured or semi-structured diagnostic interviews as the gold-standard method to establish case and control status. EHR data, on the other hand, are limited to information (e.g. billing codes, medication lists, narrative notes) collected in the course of clinical care rather than for research purposes. Recognizing this, we have undertaken systematic efforts to evaluate the validity of our EHR-based phenotyping algorithms.

In an earlier report^10^, we described the development of our automated phenotyping algorithms for BD cases and controls. Briefly, we developed four case definitions, one of which included natural language processing of narrative EHR notes and three based on structured coded data using rule-based classifiers that differed in their stringency. Another rule-based algorithm was developed to identify controls. To establish the clinical validity of these algorithms, we conducted an in-person diagnostic validation study (N = 190) in which algorithm diagnoses were compared to diagnoses made by blinded expert clinicians using a gold-standard in-person diagnostic interview (SCID-IV). Three of the four case definitions achieved high positive predictive value (PPV) compared with diagnostic interviews (up to 0.86) and the PPV for the control algorithm was 1.0. Thus, we demonstrated that automated EHR-based phenotyping can be used to identify clinically-valid case and control definitions for BD. However, an important remaining question is whether these case and control sets are *genetically* comparable to traditionally-ascertained samples that have been used in most genomic studies of BD. This is an important issue in evaluating whether EHR-based samples can be combined (e.g. through meta-analyses) with data from other ongoing genomic studies (e.g. by consortia such as the Psychiatric Genomics Consortium) to enhance gene discovery.

Here, we report genetic validation of our EHR phenotyping algorithms by using genomewide data to estimate their SNP-based heritability (h^2^_g_) and genetic correlation (r_g_) with other large-scale traditionally-ascertained BD GWAS samples. We further examined genetic correlations with other phenotypes of interest and performed genome-wide heterogeneity testing to validate the consistency of genome-wide association results. Our results demonstrate that automated EHR phenotyping can be used to assemble case/control cohorts that are both clinically and genetically comparable to traditionally-ascertained samples and thus represent a valuable tool for accelerating psychiatric genetic research.

## Materials and Methods

### Study subjects

Cases and controls were collected as part of the International Cohort Collection for Bipolar Disorder (ICCBD), a US, UK, and Swedish consortium established to accelerate genomic studies of BD by applying high throughput phenotyping methods.^7,10^ The Massachusetts General Hospital site of the ICCBD aimed to collect DNA from 4,500 cases and 4,500 controls by linking discarded blood samples to de-identified EHR data. As described in detail elsewhere^10^, cases and controls were identified by deriving EHR-based phenotyping algorithms applied to the Partners Healthcare Research Patient Data Registry (RPDR), which spans more than 20 years of data from 4.6 million patients. We first created a “datamart” of 52,235 individuals by filtering medical records to identify patients seen at Massachusetts General Hospital, Brigham and Women’s Hospital, or McLean Hospital who had at least one diagnosis of bipolar disorder (ICD-9 and DSM-IV-TR codes 296.4*–296.8*) or manic disorder (ICD 296.0*–296.1*). Next, four phenotyping algorithms were developed to identify cases and one algorithm to identify controls.

The development and clinical validation of case and control algorithms described here is adapted from Castro et al. 2015.^10^ The five phenotyping algorithms developed comprised the following:

1. 95-NLP: This BD case algorithm incorporated natural language processing (NLP) of narrative notes using the i2b2 suite of software.^11^ Expert clinicians manually reviewed 612 notes from 209 randomly selected patients to identify gold-standard cases and to extract relevant features from narrative notes to be processed by NLP. We trained a model based on 414 features to predict the probability of BD using a logistic regression classifier with the adaptive least absolute shrinkage and selection operator (LASSO) procedure. The final model, comprising 13 features, achieved an area under the receiver operating curve (AUC) of 0.93, with a sensitivity of 0.53 when the specificity was set to 0.95.
2. Coded-strict: This algorithm was a rule-based classifier that required at least three ICD codes for BD, a predominance of BD diagnoses in the longitudinal record, and either a) treatment with lithium or valproate within a year of BD diagnosis or b) treatment at a bipolar specialty clinic.
3. Coded-broad: This algorithm required at least two ICD codes for BD, a predominance of BD diagnoses, and treatment with at least two bipolar medications (lithium, valproate, carbamazepine, or an atypical antipsychotic).
4. Coded-broad-SV: This algorithm was the same as “Coded-broad” except that two or more BD diagnoses were allowed at occur during the same inpatient or outpatient episode of illness.
5. Controls: This algorithm defined controls as those age 30 years or older with no ICD-9 codes or history of medications related to a psychiatric or neurological condition.

As reported earlier, we conducted a direct-interview study to examine the predictive validity of these algorithms. Patients in the Partners Healthcare system who were identified by each algorithm as BD cases or controls were invited by mail to participate. After informed consent was obtained, participants underwent semistructured diagnostic interviews (SCID-IV) conducted by experienced doctoral-level clinicians blinded to classifier diagnosis. To further preserve clinician blinding, we recruited individuals from MGH clinics who reported a previous diagnosis of schizophrenia or major depressive disorder, disorders commonly considered in the differential diagnosis of BD. A total of 190 participants were interviewed and PPVs for each algorithm were calculated as the proportion of algorithm defined BD cases (or controls) who received a concordant diagnosis by SCID interview. The PPVs for each algorithm using a non-hierarchical approach (where each case was assigned to any algorithm for which they satisfied inclusion criteria) are shown in Table 1 and reported in Castro et al. 2015.^10^

**Table 1.** SNP-based heritability (h^2^_g_) for EHR-based bipolar disorder from the Partners Healthcare Research Patient Data Registry

| Bipolar disorder Algorithms | $h^2_g$ (SE) | | P-value<sup>2</sup> | PPV | Sample size | |
| --- | --- | --- | --- | --- | --- | --- |
|  | liability scale | observed scale |  |  | cases | controls |
| 95-NLP | 0.24 (0.10) | 0.25 (0.10) | 0.015 | 0.86 | 862 | 3952 |
| Coded-strict | 0.09 (0.05) | 0.15 (0.08) | 0.064 | 0.84 | 1968 | 3952 |
| Coded-broad | 0.13 (0.04) | 0.22 (0.08) | 0.003 | 0.80 | 2581 | 3952 |
| Coded-broad-SV | 0.00 (0.11) | 0.00 (0.18) | 0.591 | 0.50 | 408 | 3952 |
| All algorithms | 0.11 (0.04) | 0.20 (0.07) | 0.006 | NA | 3330 | 3952 |
| All algorithms except coded-broad-SV | 0.12 (0.04) | 0.21 (0.07) | 0.004 | NA | 3013 | 3952 |
| ICCBD+PGCBD<sup>1</sup> | 0.23 (0.01) | 0.41 (0.02) | $3.17 \times 10^{-80}$ | NA | 13902 | 19279 |

SNP-based heritability on liability scale was converted from observed scale based on population prevalence of 1%. <sup>1</sup>ICCBD+PGCBD: Bipolar disorder genome-wide association study from the ICCBD and PGC1 with cases ascertained by traditional methods (Charney et al. 2017). <sup>2</sup>Test for different from 0. PPV: positive predictive values from clinical validation (Castro et al. 2015). 95-NLP: probabilistic algorithm with 95% specificity based on natural language processing. Coded-strict, Coded-broad, Coded-broad-SV: coded rule-based algorithms with decreasing stringency. SV: single visit. SE: standard error.

### DNA sample collection and genotyping

The phenotyping algorithms were applied to the Partners Healthcare system to ascertain case and control DNA samples by linking phenotypic data to discarded blood samples as previously described.^11^ In brief, case and control medical record numbers are submitted to the Partners HealthCare Crimson system, which acts as an “honest broker” to match deidentified phenotypic data to discarded blood samples. Genotyping was performed in five batches that included case and control samples using the Illumina PsychChip at the Broad Institute of Harvard and MIT.

### Genotype quality control (QC) and imputation

A total of 3,772 BD cases and 4,141 controls with genomewide data were available for this analysis. We performed QC on each genotyping batch separately as follows: we removed single nucleotide polymorphisms (SNPs) with genotype missing rate > 0.05; excluded samples with genotype missing rate > 0.02, absolute value of heterozygosity > 0.2, or failed sex checks; removed SNPs with missing rate > 0.02 or with differential missing rate between cases and controls > 0.02; and removed SNPs failed Hardy-Weinberg equilibrium test (p-value < 1.0×10^−6^ in controls and p-value < 1.0x10^−10^ in cases). To merge genotyping batches for imputation and analyses, we performed batch QC by removing SNPs with differential missing rate > 0.005 between batches or significant batch association (p-value < 5.0x10^−8^ between controls form different batches). All QC were conducted using PLINK v1.9.^12^

The BD cases and controls included individuals from diverse populations. To control for population stratification and ensure the comparability between the current sample and previous European ancestry BD GWAS, we extracted samples with European ancestry for imputation and analyses. We used HapMap3 samples as a population reference panel and performed principal component analysis (PCA) with the study samples and HapMap3 samples combined. We calculated the distance between each study sample and the average European population samples in HapMap3 using PC1 and PC2. We selected the study samples with distance to average European HapMap3 samples < 0.01 (Supplementary Figure 1-3).^13^ We also removed one sample from each pair of related or duplicate samples 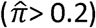.

The final analytic dataset comprised 3330 BD cases (862 95-NLP, 1968 coded-strict, 2581 coded-broad, and 408 coded-broad-SV) and 3952 controls. The sum of the individual cases groups exceeds 3330 due to the non-hierarchical design in which cases were assigned to each phenotype for which they met inclusion criteria. We performed 2-step genotype imputation with Eagle2 software for pre-phasing and IMPUTE2 on the European population study samples.^14,15^

### Statistical analysis

To assess whether our EHR-based phenotypes capture heritable components of BD, we used LD score regression (LDSC)^7,16,17^ to estimate SNP-based heritability (h^2^_g_) for each EHR-based BD cohort. We then examined the degree to which heritable influences on our BD phenotypes overlap with those traditionally-ascertained BD cases in other large-scale GWAS samples. To do this, we used LDSC to compute the genetic correlation (r^g^) between EHR-based BD samples and previously published BD GWAS by other ICCBD cohorts and the PGC (ICCBD+PGCBD).^7,16,17^ The LDSC requires association summary statistics for genome-wide SNPs to estimate h^2^_g_ and r_g_. To obtain these summary statistics, we first performed GWAS for each of the four EHR-based BD definitions separately and for our combined BD case-control sample. We used a BD prevalence of 1% to obtain liability-scale h^2^_g_ from LDSC.^18-21^ Prior studies have documented substantial genetic correlation between BD and other psychiatric disorder phenotypes, most notably schizophrenia (SCZ) and major depressive disorder (MDD).^19^ To examine the genetic relationship between EHR-based BD samples and related phenotypes, we used LD Hub^22^ to estimate r_g_ with schizophrenia (SCZ), major depressive disorder (MDD), subjective well-being, and, as a negative control, mean platelet volume (MPV). Finally, we performed genome-wide Cochran’s Q test to look for heterogeneity between association summary statistics from the EHR-based BD samples and the ICCBD+PGCBD samples at single variant level, using SNPs with association p-value < 0.001 in the ICCBD+PGCBD GWAS.

## Results

We first estimated SNP-based heritability (h^2^_g_) for the four EHR-based BD samples (Table 1). The liability-scale h^2^_g_ estimates were largest for the 95-NLP BD algorithm (0.24, p = 0.015) and smallest for the coded-broad-SV algorithm (0.0, p = 0.59), with intermediate but statistically significant values for the coded-strict and coded-broad algorithms. The h^2^_g_ of BD in the ICCBD+PGCBD sample was 0.23, which matches the h^2^_g_ for the 95-NLP algorithm but is greater than that of the rule-based algorithms. Of note, the coded-broad-SV case set had the least power with only 408 cases. As shown in Table 1, this distribution of heritability estimates mirrors the relative PPVs obtained in our clinical validation study. To maximize the BD case-control sample size, we combined the BD case-control samples across algorithms into a single case-control dataset. Since the coded-broad-SV had no evidence of heritability, we created two combined BD datasets; one included all BD cases and one included all but the coded-broad-SV cases). The h^2^_g_ was 0.11 (p-value = 0.006) for all algorithms combined BD and 0.12 (p-value = 0.004) for all algorithms excluding coded-broad-SV.

We next estimated the SNP-based genetic correlation (r_g_) between the EHR-based BD samples and the ICCBD+PGCBD samples (Table 2). The r_g_ estimates were 95-NLP (0.66), coded-strict (1.0), and coded-broad (0.74) were all statistically significant. (Note that r_g_ could not be estimated for coded-broad-SV given its h^2^ _g_ of 0). The r_g_ for all algorithms excluding coded-broad-SV was 0.83 (p= 7.19x10^−7^). Adding coded-broad-SV BD cases to the combined case set did not substantially change the (r_g_) estimate although the standard error (SE) increased and p-value rose to 2.88x10^−6^. We also estimated the pairwise r_g_ between the EHR-based BD case-control samples and the final combined BD samples (Table 3). The r_g_ estimates ranged from 0.90 to 0.98 between algorithms, and were 1.00 between each algorithm and the combined sample (excluding coded-broad-SV). Finally, the r_g_ between ICCBD and PGCBD was 1.00 (SE = 0.065, p-value = 1.45x10^−74^).

**Table 2.** SNP-based genetic correlation (r_g_)between EHR-based bipolar disorder and bipolar disorder ascertained by traditional methods from ICCBD+PGCBD

| | $r_g$ (SE) | P-value<sup>1</sup> |
| --- | --- | --- |
| 95-NLP | 0.66 (0.16) | $3.69 \times 10^{-5}$ |
| Coded-strict | 1.00 (0.29) | $2.40 \times 10^{-4}$ |
| Coded-broad | 0.74 (0.15) | $8.11 \times 10^{-7}$ |
| All algorithms | 0.83 (0.18) | $2.88 \times 10^{-6}$ |
| All algorithms except<br>Coded-broad-SV | 0.83 (0.17) | $7.19 \times 10^{-7}$ |

Genetic correlation was not estimated for Coded-broad-SV due to SNP-based heritability estimate of 0. Genetic correlation ( $r_g$ ) was estimated against PGC bipolar disorder GWAS. <sup>1</sup>Test for different from 0. SE: standard error.

**Table 3.** SNP-based genetic correlation (r_g_) between EHR-based bipolar disorder

| Phenotype 1 | Phenotype 2 | $r_g$ (SE) | P-value<sup>1</sup> |
| --- | --- | --- | --- |
| 95-NLP | Coded-strict | 0.90 (0.19) | $1.32 \times 10^{-6}$ |
| 95-NLP | Coded-broad | 0.96 (0.13) | $3.65 \times 10^{-13}$ |
| Coded-strict | Coded-broad | 0.98 (0.08) | $1.28 \times 10^{-34}$ |
| All algorithms except coded-broad-SV | 95-NLP | 1.00 (0.12) | $1.05 \times 10^{-16}$ |
| All algorithms except coded-broad-SV | Coded-strict | 1.00 (0.07) | $3.34 \times 10^{-54}$ |
| All algorithms except coded-broad-SV | Coded-broad | 1.00 (0.01) | $1.91 \times 10^{-991}$ |

<sup>1</sup>Test for different from 0. SE: standard error.

Given prior evidence that traditionally-ascertained BD GWAS show significant positive genetic correlations with SCZ and MDD^17,19^ and significant negative genetic correlation with subjective well-being^23^, we examined these correlations using our EHR-based algorithms as another index of their genetic validity. As a negative control, we also examined their genetic correlation with mean platelet volume, a phenotype for which we would not expect significant genetic correlation. (Figure 1; Supplementary Table 1). We used the cross-phenotype r_g_ of ICCBD+PGCBD as the standard for comparison. As expected based on prior data^17,19,23^, the EHR-based case-control samples positively correlated with SCZ and BD, negatively correlated with subjective well-being, and uncorrelated with MVP (Figure 1). These patterns were mirrored those observed for the ICCBD+PGCBD sample with one difference. Whereas the genetic correlation was greater between EHR-based BD and MDD was larger than that seen for EHR-based BD and SCZ, the opposite order was seen between ICCBD+PGCBD and these phenotypes. This difference in magnitude remained when r_g_ were estimated separately for ICCBD and PGCBD.

**Figure 1:**
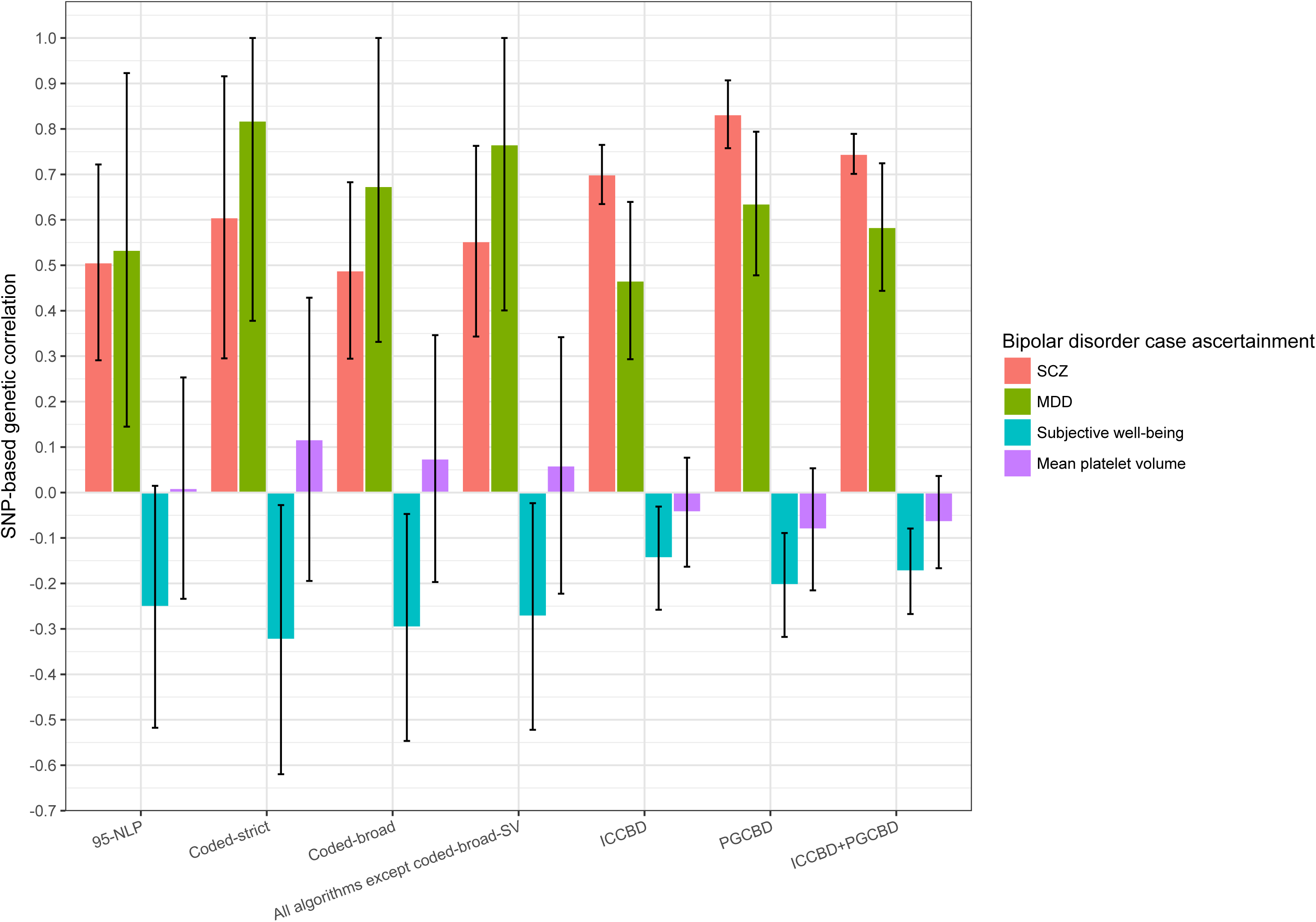
SNP-based genetic correlation (with 95% confidence interval) between bipolar disorder based on different ascertainment methods and other traits

We hypothesized that this difference in r_g_ patterns might be related to differences in the proportions of BD subtypes among the EHR-BD cases and those included in the traditionally-ascertained samples. To investigate this, we calculated the percentage of BD case subtypes, including bipolar I disorder (BD1), bipolar II disorder (BD2), schizoaffective disorder bipolar type (SAB), and bipolar disorder not otherwise specified (NOS) for the EHR-based BD cases and the ICCBD cases (the subtypes of PGCBD cases were not available). We found that the EHR-based BD cases comprised a lower proportion of SAB subtype cases (0.6-1.6%) compared with the ICCBD samples (9.1%) (Supplementary Figure 4). This difference would be consistent with a relatively larger genetic correlation with SCZ seen with the ICCBD sample compared to the EHR-based samples.

Finally, we performed Cochran’s Q test to identify potential heterogeneity of the association summary statistics between EHR-based BD samples and the ICCBD+PGCBD samples. This analysis was restricted to SNPs with association p < 0.001 in the ICCBD+PGCBD GWAS in order to exclude SNPs with weak association results whose directionality might be less robust. We identified a single locus with significant heterogeneity across the genome after Bonferroni correction (SNP N = 28,320) for both coded-broad and for the combined EHR-BD sample (excluding coded-broad-SV) (Figure 2). This locus on chromosome 22 (peak Q test p-value at rs196065 = 3.34x10^−7^), showed modest association with BD (p-value = 5.78x10^−5^ in ICCBD+PGCBD) and did not overlap with any previously reported BD-associated loci. Thus, we found negligible evidence of heterogeneity of genomewide association results between EHR-based BD and traditionally-ascertained BD.

**Figure 2:**
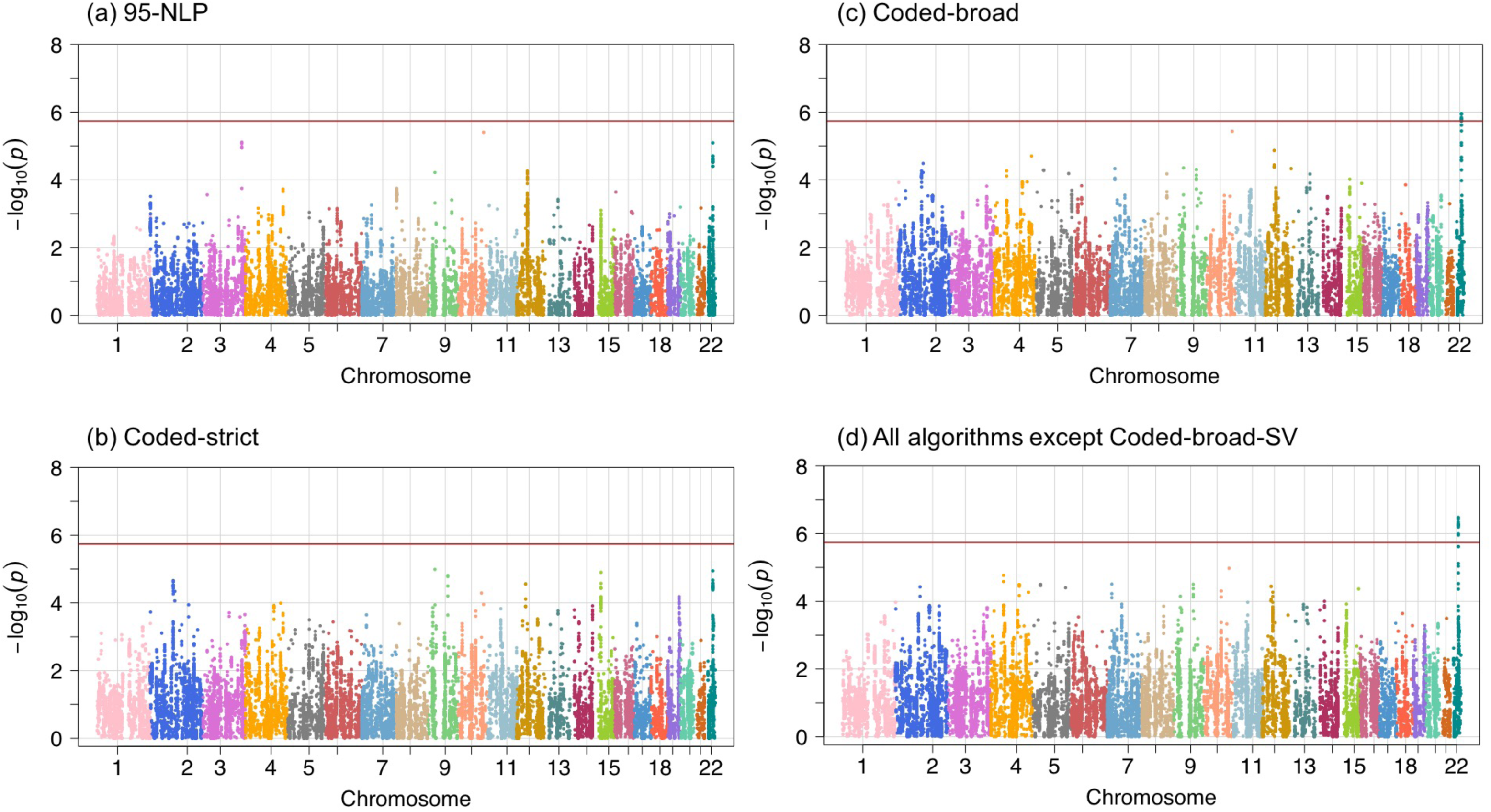
Genome-wide Cochran’s Q-test for heterogeneity of SNP effects between ICCBD+PGCBD and EHR-based bipolar disorder. Red line shows the Bonferroni-corrected significance level for the Q-test. SNPs are selected with association p-value threshold of 0.001 based on ICCBD+PGCBD analysis (total number of SNPs=28,320).

## Discussion

As an ever-growing longitudinal repository of the clinical phenome, EHRs represent a new and powerful resource for psychiatric research.^9^ Nevertheless, their utility depends on the validity of the clinical and phenotypic data that can be extracted. We have previously demonstrated the feasibility of deriving diagnoses with high predictive value compared with a gold standard of clinician-administered diagnostic interviews.^10^ However, in the context of psychiatric genetic research, establishing the genetic validity of these phenotypes is crucial. In the present study, using genomewide genotype data for more than 7,000 cases and controls, we demonstrate that EHR-based algorithms can be used to ascertain BD phenotypes that are heritable and genetically comparable to traditionally-ascertained samples. Automated algorithm-based phenotyping linked to biospecimens provides substantial efficiencies in terms of the time and costs involved in assembling large-scale samples for genetic research. Prior simulations have documented up to a 10-fold reduction in the cost associated with phenotyping and sample collection.^11^ Using our case/control BD definitions linked to discarded blood samples, we were able to collect approximately 5,000 controls over 10 weeks and more than 4,000 cases over 3 years. As described below, three sets of findings from our analyses are particularly noteworthy.

First, our results document that EHR-based diagnostic algorithms can be used to ascertain BD phenotypes that yield SNP-based heritability comparable to that observed in GWAS that have relied on more time-cost-, and labor-intensive recruitment and diagnostic evaluation. The highest heritability (0.023) was seen with our 95-NLP algorithm which combined NLP of narrative test features and coded EHR data. This estimated heritability was nearly identical to that derived from GWAS of the larger traditionally-ascertained cohorts of the international ICCBD and PGC (h^2^_g_ =0.24 for 13,902 cases and 19,279 controls). The 95-NLP algorithm also achieved the highest positive predictive value in our previous clinical validation study. For two of the remaining three algorithms which involved rule-based algorithms of structured EHR data, we also observed significant, though relatively lower, heritability estimates (h^2^_g_ = 0.09 – 0.12). The least restrictive algorithm (coded-broad-SV) did not exhibit significant heritability, though the small sample size of this subgroup may limited the power of our analyses. Of note, this last algorithm also performed poorly in our prior clinical validation study (PPV=0.5). Nevertheless, the overall heritability of our EHR-based BD was 0.12 (p = 0.004), dropping slightly to 0.11 (p = 0.006) when the coded-broad-SV was included. In addition, the EHR-based BD definitions were nearly perfectly genetically correlated. Pairwise genetic correlations between the phenotypes ranged from 0.98 – 1.0 except for 95-NLP and coded-broad-SV (r_g_= 0.90).

Second, we found that our cohorts ascertained by automated EHR phenotyping exhibited substantial genetic correlations (r_g_) with the large ICCBD+PGCBD samples. Overall, the r_g_ between our EHR-based BD case/control samples and the ICCBD+PGCBD samples was 0.83 (p = 2.88 x 10^−6^), demonstrating that our approach captures genetic influences that strongly overlap with those acting on BD in traditionally-ascertained samples. In addition to providing further genetic validation of EHR-derived phenotypes, these results indicate that such samples can be combined with other existing samples to enhance the power of genetic discovery.

Finally, we demonstrate that our phenotyping approach replicates patterns of cross-disorder genetic overlap that have previously been reported in genetic studies of BD.^7,24^ In particular, EHR-based BD exhibited positive genetic correlations with SCZ and MDD and negative correlations with subjective well-being. Once again, this supports the genetic validity of our algorithm-defined BD phenotype. Unexpectedly, the genetic correlation with SCZ was less than that seen with MDD, a finding that may be attributable to the relatively low frequency of SAB cases in our sample.

We acknowledge that our results have certain limitations. First, our sample size, while substantial, is smaller than that of some other existing samples (e.g. ICCBD and PGCBD), which may have limited the power and precision of our heritability and genetic correlation analyses. Second, the portability of our specific phenotyping algorithms to other healthcare settings remains to be determined. Notably, however, our results demonstrate that a range of algorithms – with and without NLP and using diagnostic rules of varying stringency – yield phenotypes that are clinically and genetically comparable to those obtained by in-person standardized diagnostic assessments.

In summary, the current study provides the first genetic validation of EHR-based phenotyping for BD and suggests that automated phenotyping algorithms can identify samples that are highly genetically correlated with those ascertained through conventional methods. Taken together, the present results and those of our prior clinical validation study, suggest that the use of any or all three of the heritable EHR-based algorithms we derived (i.e. 95-NLP, coded-strict, and coded-broad) can facilitate genetic studies of bipolar disorder. High throughput phenotyping using the large data resources available in the EHR database represents a viable method for accelerating psychiatric genetic research.

## Acknowledgments

This work was supported in part by NIMH grants R01MH085542 (JWS and PS), R01MH085545 (JWS), and K24MH094614 (JWS) and by support from the Demarest Lloyd, Jr. Foundation. Dr. Smoller is a Tepper Family MGH Research Scholar.

## Disclosures

Dr. Smoller is an unpaid member of the Scientific Advisory Board of PsyBrain Inc. and the Bipolar/Depression Research Community Advisory Panel of 23andMe.

